# Flow augmentation from off-channel storage improves salmonid habitat and survival

**DOI:** 10.1101/2023.02.15.528508

**Authors:** Gabriel J. Rossi, Mariska Obedzinski, Shelley Pneh, Sarah Nossaman Pierce, William T. Boucher, Weston M. Slaughter, Keane M. Flynn, Theodore E. Grantham

## Abstract

In the Western United States, juvenile salmon and steelhead are especially vulnerable to streamflow depletion in the dry season. Releasing water from off-channel storage is a method of streamflow augmentation increasingly used to offset impacts of anthropogenic flow alteration. However, to date, no studies have evaluated the effects of these small-scale flow augmentations on salmonids. Here we quantify the effects of one such augmentation project on habitat connectivity, water quality, invertebrate drift, juvenile salmonid movement and survival. Our study took place in a Northern California stream and included an unusually wet summer (2019) and a more typical dry summer (2020). We found that differences in ambient streamflows between the two years mediated the physical and ecological effects of a 13.9 L/s augmentation treatment. In the dry year, flow augmentation significantly improved dissolved oxygen and habitat connectivity at sites > 1.5 km downstream from the point of augmentation and had a marginal warming effect on stream temperature. During the wet year, both dissolved oxygen and water temperature effects were negligible. In both years, augmentation had a small but positive effect on invertebrate drift. Inter-pool movement of juvenile steelhead (Oncorhynchus mykiss) and stocked Coho Salmon (O. kisutch) increased due to augmentation during the dry summer. Flow augmentation also increased the survival probability for salmonids, with a larger effect during the dry summer (24% higher survival for Coho and 20% higher for steelhead), than during the wet summer (when no effect was observed for steelhead survival and Coho Salmon survival increased by 11%). This study indicates that appropriately designed and timed flow augmentations can improve conditions for rearing salmonids in small streams, particularly during dry years. More broadly it provides empirical evidence that efforts to restore summer streamflow in small, salmon-bearing streams can yield significant ecological benefits.

## [A] Introduction

Streamflow alteration is ubiquitous in the United States (Carlisle et al. 2019) and has been identified as a primary stressor to imperiled Pacific salmon and trout (*Oncorhynchus* spp.) populations in the Western US (Moyle et al. 2017; Crozier et al. 2019). In California, salmonids are particularly vulnerable to flow depletion in the dry season—a period of naturally low flow in which stream habitats contract, water quality conditions deteriorate, and food resources become limited (Vander Vorste et al. 2020, Obedzinski et al. 2018, Caldwell et al. 2018). A growing body of research from California has shown that for rearing salmonids in many streams, the dry season creates a metabolic knife’s edge – where only a few days of hydraulic disconnection, a slight change in dissolved oxygen levels, or few degrees of temperature separates profitable from unfavorable habitat conditions (Harvey et al. 2006; Boughton et al. 2009; Grantham et al. 2012; Obedzinski et al. 2018; Hwan et al. 2018; Sloat and Osterback 2012; Woelfle-Erskine et al. 2017, Vander Vorste et al. 2020). Water diversions during the dry season can compound these natural stressors, tipping the balance from profitable to stressful or even lethal environments for rearing juvenile salmonids (Deitch et al. 2009, Grantham et al. 2012, Power et al. 2015, Obedzinski et al. 2018). This study explores whether relatively small augmentation of flow in the dry-season flow can have a reciprocal effect and improve habitat, feeding opportunity, and survival of salmonids.

Recently, streamflow augmentation from off-channel storage has been implemented opportunistically as a restoration strategy for salmonids in Northern California (Deitch and Dolman 2017, Ruiz et al. 2019, Russian River Coho Partnership 2020). Rather than using recycled water to augment flow in urban streams (Halaburka et al. 2013), these projects use surplus water from agricultural irrigation ponds or storage tanks specifically to enhance flow in salmon-bearing streams in rural watersheds (Ruiz et al. 2019, Russian River Coho Partnership 2020). While initial data suggests this type of flow augmentation can improve summer habitat conditions for rearing salmonids (Ruiz et al. 2019, Russian River Coho Partnership 2020), no experimental studies have been conducted to quantify the effects of flow augmentation on juvenile salmonid growth and survival in these small, undammed streams. Here we evaluate data from a streamflow augmentation study on Porter Creek, a small, salmon-bearing tributary to the Russian River in Northern California. This study specifically investigated the response of summer-rearing salmonids, and the habitat they depend on, to changes in streamflow. Our objectives were (1) to evaluate the spatial effects of flow augmentation on habitat connectivity and (2) to measure the response of physical and biotic variables that influence the rearing profitability and fitness of summer-rearing salmonids to a controlled flow augmentation treatment.

The flow augmentation experiment released of 13.9 L/s of water from an off-stream pond into the stream channel for one month in the summers of 2019 and 2020. Augmentation was initiated as the first riffles began to dry. To address the first objective, we measured the extent of wetted channel from the point of augmentation (2.75 km) downstream to the creek outlet. Wetted channel extent below the augmentation release point was compared to an unaltered, control reach located approximately 2 km upstream of the augmentation (Figure 1). To address the second objective, we used a multiple before–after-control-impact time series (mBACIPs) statistical analysis (Keough and Mapstone, 1997, Wauchope et al. 2020), to estimate how flow augmentation affected dissolved oxygen, water temperature, hydraulic habitat, invertebrate drift, and inter-pool movement of fish. We also estimated the effects of flow augmentation on the survival of juvenile steelhead (*Oncorhynchus mykiss)* and Coho Salmon (*O. kisutch)* using a mark-recapture approach and robust design model implemented by Obedzinski et al. (2018). We hypothesized that (H1) augmentation would increase streamflow and hydraulic connectivity, and that (H2) the effects of augmentation would diminish over space and time (distance downstream from the point of augmentation and as the dry season progressed). We also hypothesized that the augmentation treatments would: (H3) increase dissolved oxygen by decreasing flow residence time, (H4) increase water temperature if augmentation was warmer than ambient flow (and vice versa), and (H5) increase invertebrate drift. Finally, we hypothesized that augmentation would increase (H6) salmonid movement, (H7) growth, and (H8) survival relative to the non-augmented control reach.

**Figure 1.**
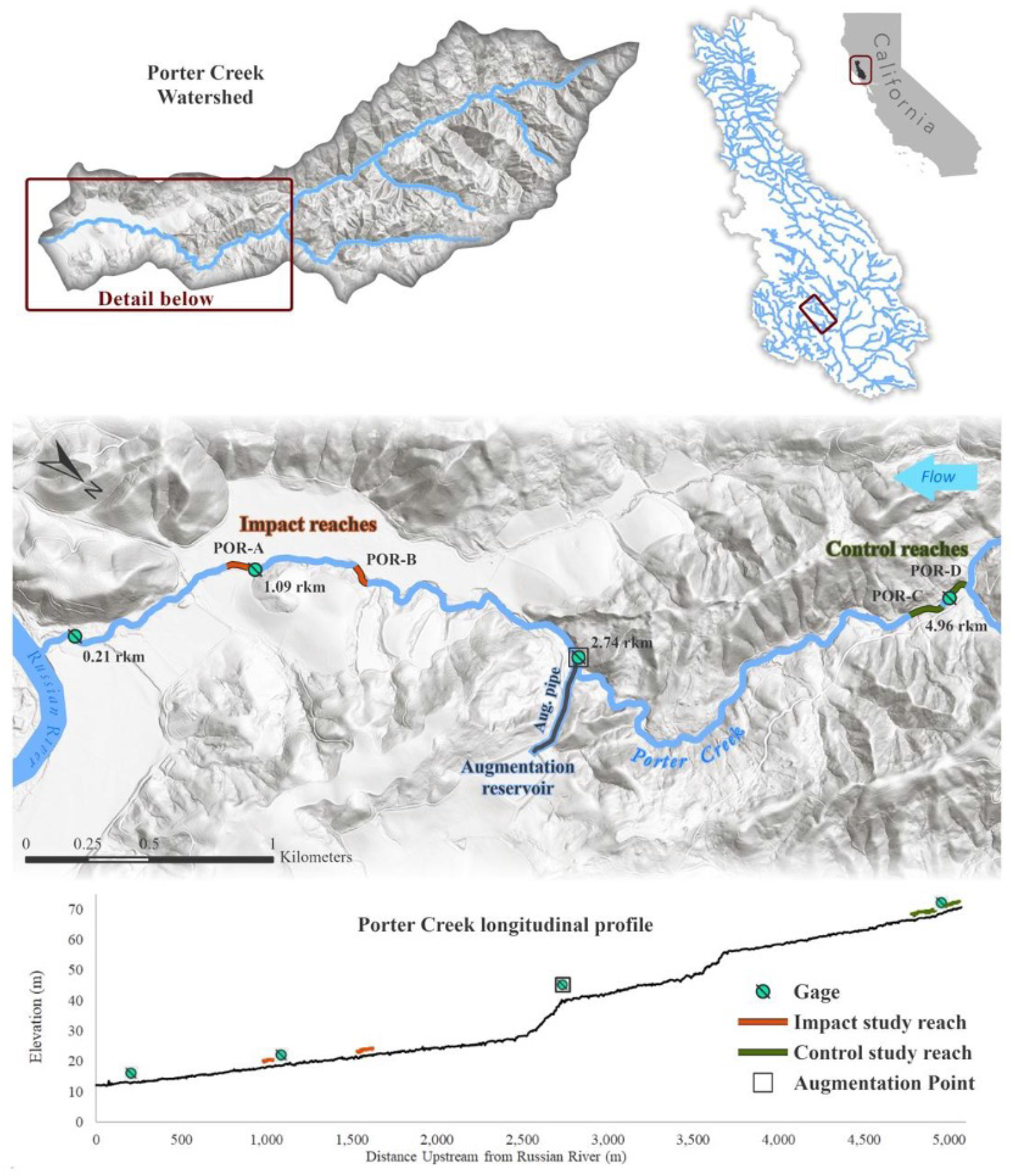
Porter Creek watershed (top left), located in the Russian River watershed in Northern California (top right). The study area map of lower Porter Creek (middle) shows flow gaging sites, control and impact study reaches, the off-channel pond, and the augmentation pipe (dark blue line). The longitudinal profile (bottom) shows the relative location of gaging stations, the augmentation point, and the control and impact study reaches.

## [A] Methods

### [B] Study location and project history

Porter Creek is a tributary to the Russian River in Sonoma County, California (Figure 1). The 19.4 km^2^ watershed is located in California’s Coast Range and flows approximately 11.4 km west to east into the Russian River, near the town of Healdsburg. The watershed is entirely privately owned and managed for timber, livestock, and premium wine-grape production, with extensive vineyard planting on terraces in the lower 2.3 km of the stream valley. Rural residential development occurs in the upper watershed. The watershed’s geology is primarily composed of Franciscan complex mélange, although the southern slopes and upper watershed are a mix of mélange and coastal belt shales (Jennings 1977, California Geological Survey, Geologic Data Map No. 2). The Franciscan mélange geology is regionally associated with sparse deciduous oak and annual grass savanna and limited subsurface storage capacity to sustain streamflows throughout the dry, warm summer season (Hahm et al. 2019).

Streamflow patterns in Porter Creek are characterized by Mediterranean hydrology. More than 90% of the annual rainfall typically occurs from November through April, resulting in flashy winter flows and then steady streamflow recession following the last spring freshets until flows cease completely pending the first fall rains (Deitch et al. 2009). Except in the wettest years, Porter Creek becomes intermittent for much of its length by mid- or late-summer. As flows recede, riffles dry first, leaving isolated pools, followed by the contraction of wetted pool habitat and, in some stream reaches, complete channel drying by late-summer.

Porter Creek supports both wild and conservation-hatchery-stocked Coho Salmon (*Oncorhynchus kisutch)*, wild steelhead (*Oncorhynchus mykiss*), and a number of native cyprinid fishes – most commonly California Roach (*Hesperoleucus symmetricus*) and sculpin (*Cottus spp)*. Russian River salmonid populations, particularly Coho Salmon, have experienced rapid declines over the last century, with fewer than 10 adult Coho Salmon observed returning each year by the early 2000s. These declines led to state and federal endangered species listings (NMFS 2012) and prompted a multi-agency conservation hatchery effort designed to raise and release juvenile Coho Salmon in key Russian River tributaries (NMFS 2012, Obedzinski et al. 2018). Porter Creek was listed in the National Marine Fisheries Service Coho Salmon Recovery Plan as a priority stream for salmonid restoration and streamflow improvement (NMFS 2012) and has received planted fish from the Don Clausen Fish Hatchery at Warm Springs Dam most years since 2010.

The 2012 - 2016 California drought was the driest five-year period in California history since modern record-keeping began (Hanak et al. 2015) and precipitated several recovery actions for imperiled salmonid species. In 2014, the State of California passed the Water Quality, Supply, and Infrastructure Improvement Act of 2014, which authorized the Legislature to appropriate $200 million towards projects designed to enhance stream flows (California Legislature 2014). While flow enhancement was already underway, this (and subsequent funding acts) spurred numerous small flow enhancement projects across coastal California. In 2014, the primary vineyard landowner on Porter Creek entered into a voluntary drought agreement with state management agencies to release water stored in an off-stream reservoir into the stream channel, 2.74 km upstream of the creek outlet, to maintain suitable rearing conditions for juvenile salmon and to facilitate passage of out-migrating smolts (Figure 1). This storage reservoir is filled with water pumped from shallow wells along the Russian River, upstream of the confluence with Porter Creek, in addition to annual precipitation recharge. The reservoir elevation is above the augmentation structure and water is gravity fed into a plumbing array, through a series of butterfly valves, and downslope through a pipe to the augmentation point in Porter Creek (Figure 1, Supplemental Materials Figure S1 and S2). A programmable controller regulates flow releases from the reservoir (Supplemental Material, Supplemental Materials Figure S1) and allows an operator to set the start date and duration of specific augmentation levels. The completed streamflow augmentation system can release up to 61,000 m^3^ of stored water into the creek each year, at a rate of up to 26 L/s (0.9 ft^3^/s).

### [B] Experimental Study Design

Four study reaches were selected on Porter Creek – two in a control reaches (upstream of augmentation between river kilometer rkm 3.7 and rkm 4.65) and two in an impact reaches (downstream of augmentation between rkm 1.0 to 1.65; Figures 1). Each study reach contained four riffle-pool habitat units which served as replicates, resulting in eight “control” units and eight “impact” units (Figure 2). The riffle-pool unit is a ubiquitous geomorphic feature in alluvial streams with slopes between 0.5% and 2% (Leopold and Wolman 1957) and provides a discrete habitat unit for evaluating juvenile salmonid rearing and foraging during the low-flow period (Rossi et al. 2021). Study site selection was constrained by landowner access and channel confinement in the upper reaches of Porter Creek and was, therefore, not random. We selected reaches with similar slope and geomorphic characteristics that supported foraging salmonids and we prioritized sites with simple hydraulic controls so that we could develop robust rating curves at the downstream riffle crest.

**Figure 2.**
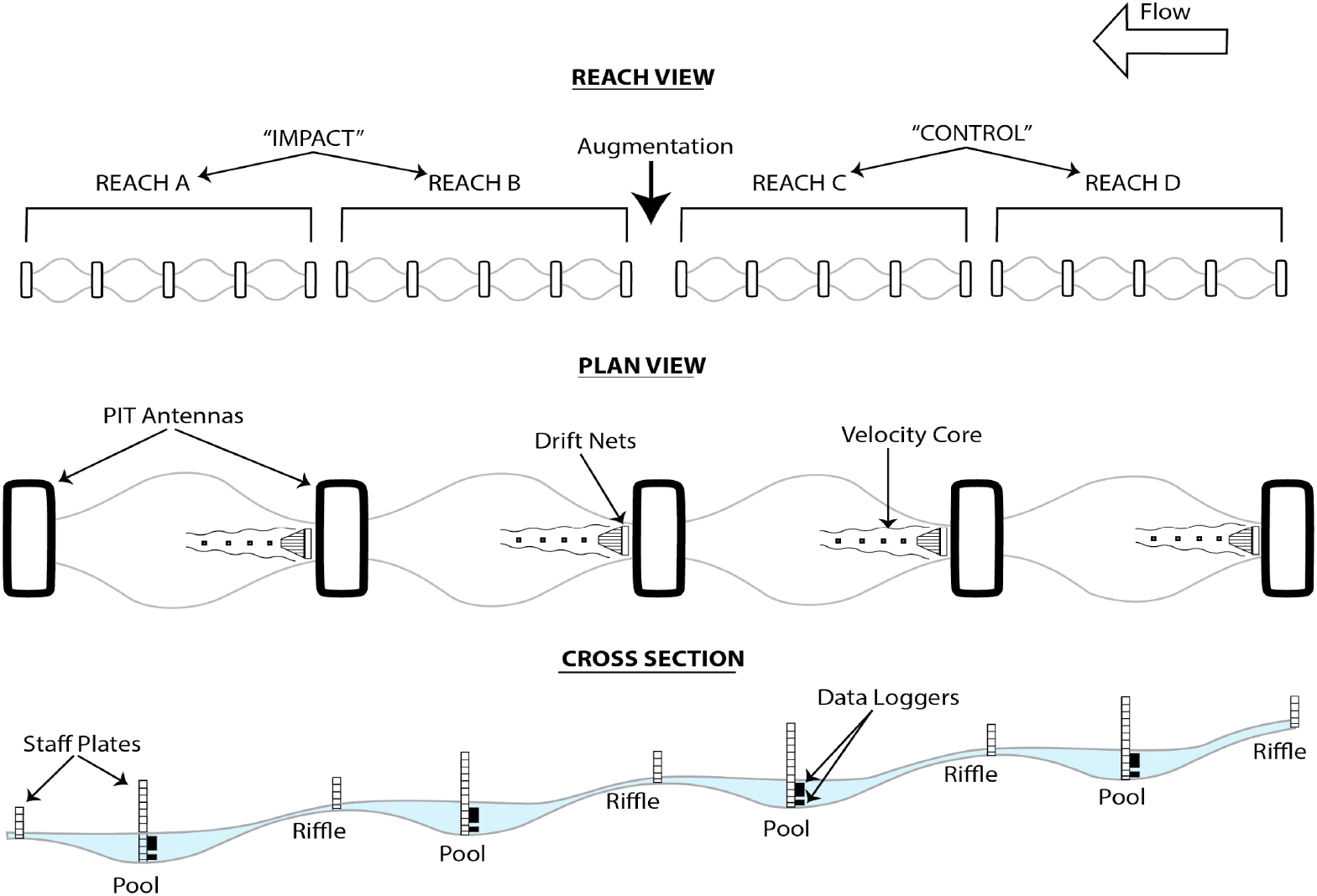
Schematic of the Porter Creek study design showing two “control” and two “impact” study reaches separated by the point of augmentation (top), as well as a plan view (middle) and cross-section view (bottom) of a study reach. Each study reach contained four consecutive riffle- pool units, bounded by a PIT antenna (see *Salmonid Movement*) and containing a staff plate, HOBO U26 DO logger and HOBO U20 water logger in the pool, and a second staff plate on the downstream riffle crest. The drift net (see *Invertebrate Sampling)* and pool velocity measurement locations are also shown.

### [B] Streamflow Augmentation

We tested the same level of flow augmentation during a wet summer (2019) and a dry summer (2020) (Figure 3). Experimental augmentations were timed to occur just prior to riffle-pool disconnection in the control reach of Porter Creek. In 2019, a large May freshet extended surface connectivity until mid-July in the control reach (Figure 3). We tested an augmentation treatment of 13.9 L/s, starting on July 12^th^ and continuing through October. This treatment (13.9 L/s) was repeated during the much drier summer of 2020, starting on June 25^th^ and ending on August 6 (Figure 3). Streamflow was gaged at three locations below augmentation (rkm 2.75, 1.05 and 0.25) and one location upstream (rkm 4) (Figure 1). In 2019, augmentation commenced when flow was 5 L/s at rkm 4 (control reach) and 25 L/s at rkm1.05. In 2020, augmentation commenced when flow was 4.2 L/s at rkm 4 and only 4.3 at rkm 1.05. The difference in downstream accretion between 2019 and 2020 was due to the increased subsurface water storage after the May freshet in 2019.

**Figure 3.**
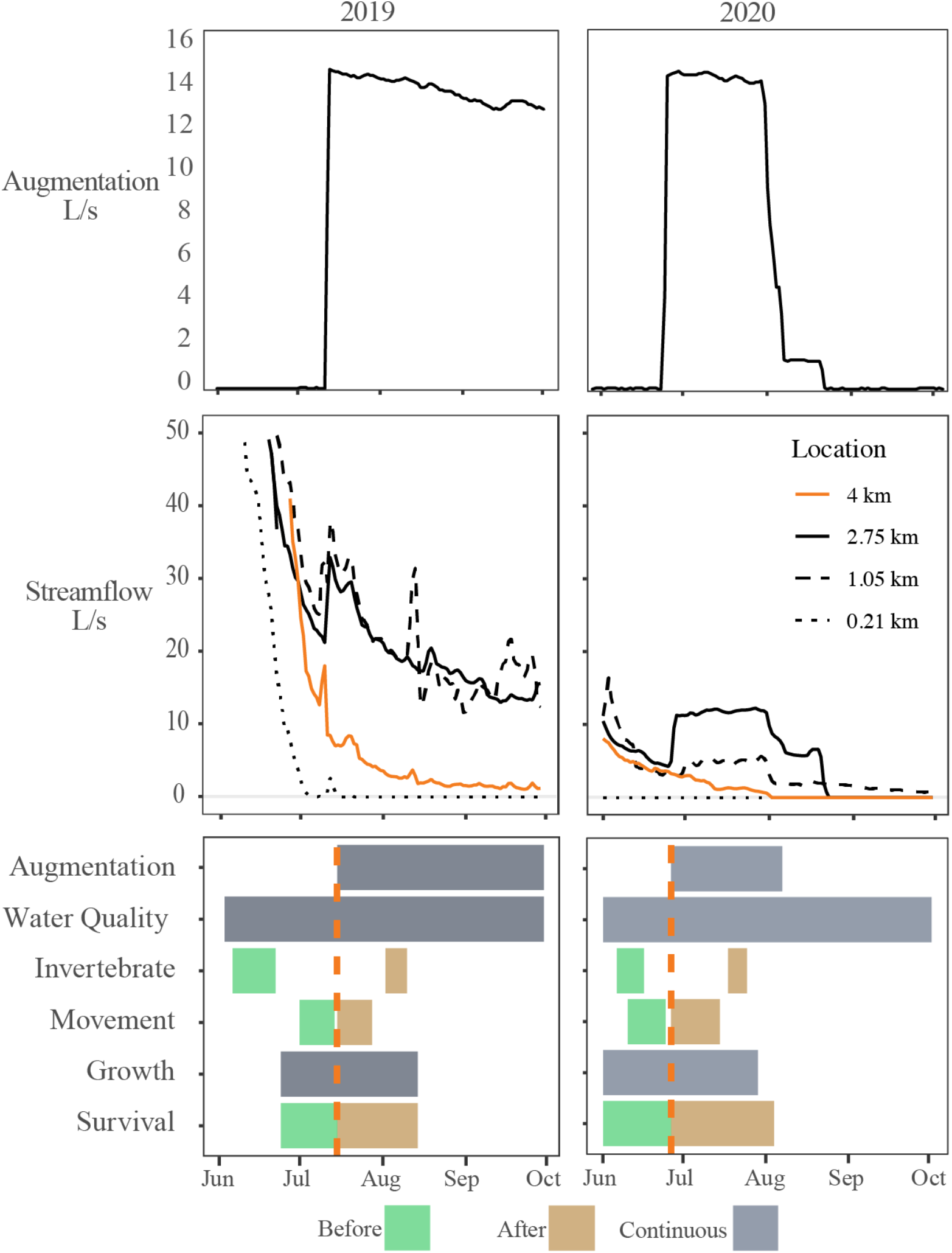
Top row: streamflow augmentation schedule, summers 2019 (left) and 2020 (right). Middle row: ambient streamflow above the augmentation (orange, 4 km), at the augmentation (black solid, 2.75 km) and downstream (dashed lines) in 2019 (left) and 2020 (right). Bottom row: the timing of augmentation relative to data collection for parameters measured in this study in 2019 (left) and 2020 (right).

### [B] Wetted Habitat Mapping

The spatial extent of “wet” (flowing stream), “intermittent” (dry riffles and wet pools), and “dry” (dry riffles and pools) channel was mapped immediately prior to and one month after augmentation. These wetted habitat surveys began at the confluence with the Russian River and continued upstream 5.01 km to the top of the control reach (Figure 1). Data were entered using the mobile ArcCollector application with a handheld GPS unit (BadElf, 2.5m stationary accuracy). The complete field protocol for the wetted habitat surveys is available online (CA Sea Grant 2019). Summary statistics describing the total proportion of wet, intermittent, and dry channel between rkm 0 to rkm 5.01 were computed for each surveyed interval and related to augmentation level and ambient streamflow.

### [B] Hydraulic Habitat Measurements

Onset HOBO U20 water level loggers were deployed in all pools within each study reach to measure continuous changes in pool depth. Water level loggers were mounted to rebar and installed near the pool maximum depth (Figure 2). Depth was measured where the thalweg bisects the downstream riffle crest of each pool, a point known as the ‘riffle crest thalweg’ (RCT) (Rossi et al. 2021). RCT depth serves as an indicator of hydraulic connectivity and has been shown to be correlated with dissolved oxygen concentrations and fish behavior (Rossi et al. 2021, Rossi et al. 2021b). The elevation of the RCT was surveyed relative to the elevation of the water level logger and staff plate to allow for a conversion between continuous pool stage and continuous RCT depth (Figure 2). Velocity was measured along the thalweg of the pool from the upstream point where water entered the pool at one-meter increments (Figure 2). These velocity profiles served as a proxy for the changing length of feeding zones in pools below riffles due to augmentation (Harvey et al. 2006).

### [B] Water Quality Monitoring

Continuous dissolved oxygen and stream temperature were measured using Onset HOBO U26 data loggers in each pool (Figure 2). The dissolved oxygen loggers were lab calibrated prior to deployment. Field calibration measurements were taken three times during the study period using a handheld YSI Pro20 in each pool and these values were used to correct the logger output data using HOBOware Pro’s Dissolved Oxygen Assistant software.

### [B] Invertebrate Sampling

To estimate the effect of streamflow augmentation on prey abundance for salmonids and the stream invertebrate assemblage we sampled benthic macroinvertebrate (BMI) drift entering each pool. Invertebrate drift was sampled two weeks before and two weeks after augmentation in both years (Figure 3). Sampling took place between 16:00 and 19:00 each day. Drift was collected in nets with a 50 cm × 20 cm mouth aperture and 500-μm mesh. The drift net was installed just below the water surface at the head of each pool (Figure 2). Cross-stream position of the net was adjusted on each occasion to capture the region of highest velocity at each sampled streamflow. All invertebrate samples were preserved in 100% ethanol in the field, and in the lab, identified to the closest taxonomic value and measured to the nearest 0.5mm with a dissecting scope (Merritt et al. 2008). Each invertebrate was measured to the nearest 0.5 mm under a dissecting scope and biomass (mg dry mass) was estimated from published length to dry mass relationships (Benke et al. 1999; Sabo et al. 2002). Since our statistical analysis was intended to evaluate the effect of invertebrate availability as a food resource for fish, we removed large (>10mm) taxa which made up less than 0.5% of the total drift but skewed the biomass estimates.

### [B] Study Population

Salmonid movement, growth, and survival were evaluated using a population of passive integrated transponder (PIT) tagged fish. As an initial marking event, Coho Salmon and steelhead juveniles were captured from each study reach (Figure 2) using backpack electrofishing during sampling events on June 24^th^ and 25^th^, 2019, and June 1^st^ and 2^nd^, 2020 (Figure 3). Fish larger than 60 mm were fitted with 12 mm PIT tags by creating a small (1mm) incision in the body cavity on the ventral side and inserting the tag (Table 1). In addition to the fish captured and tagged in the stream environment, tagged juvenile Coho Salmon raised for the Russian River Coho Salmon Captive Broodstock Program at the Don Clausen Fish Hatchery at Warm Springs Dam were stocked into reaches A and C (Figure 2) in both years. Approximately 250 Coho Salmon were stocked into each of the two reaches on June 27^th^, 2019 (mean of 65 fish/pool) and June 8^th^, 2020 (mean of 62 fish/pool) (Table 1).

**Table 1.** Counts of salmonids that were PIT tagged in Porter Creek (left) or tagged individuals released into Porter Creek from Don Clausen Fish Hatchery (right) in 2019 and 2020. Hatchery coho salmon were released only into reaches A and C (Figures 1 and 2).

| Species | 2019 |  |  | 2020 |  |  |
| --- | --- | --- | --- | --- | --- | --- |
|  | Wild | Hatchery | Total | Wild | Hatchery | Total |
| Steelhead | 186 | 0 | 186 | 174 | 0 | 174 |
| Coho Salmon | 5 | 517 | 522 | 166 | 497 | 663 |

### [B] Salmonid Movement

Movement of all PIT-tagged fish was monitored using stationary PIT antenna arrays mounted over riffles between each study pool (Figure 2). Antennas were mounted 30 cm above the water surface on cinder blocks to allow fish to swim freely while maintaining proximity to the antennas for detection. The PIT antennas were connected to power sources and a Biomark IS1001 data logger located on the adjacent terrace. Each reader was programmed to log hourly status reports which documented antenna operation to ensure they were consistently running. To estimate the effect of augmentation on salmonid movement between pools and riffles, we computed the total number of detections at each antenna per day (in the control and impact reaches), and also the number of detections per unique tag per day so that a single mobile fish would not bias the results.

### [B] Salmonid Growth

Fish growth was estimated by measuring the fork length (mm) of recaptured individuals and comparing those measurements to fork length at the time of tagging. Growth intervals were 52 days in 2019 and 57 days in 2020 (Figure 3). These intervals include time before and after augmentation was initiated, but more than half the duration of the growth intervals (33 days in 2019 and 34 days in 2020) occurred after augmentation (Figure 3). Growth was computed as change in length (mm/day). We differentiated fish smaller than 80 mm as “young-of-year” (YOY) from larger “parr” (>80 mm) but restricted our statistical analysis to YOY because too few parr were captured. No smolts were observed during our summer sampling.

### [B] Salmonid Survival

To estimate reach-specific survival before and after augmentation, we used an individual-based capture-mark-recapture approach that consisted of an initial marking event and two subsequent primary recapture occasions each year – one before the onset of augmentation and one following a period of augmentation. The initial electrofishing surveys in which the fish were tagged or the date of hatchery release served as the initial capture and marking event (Table 1). Each of the two recapture occasions consisted of two consecutive days of PIT-tag “wanding” in which biologists waded the full extent of each reach, detecting as many fish as possible using a portable PIT-tag detection system, or “wand”. This sampling approach allowed estimation of survival probability using the robust design model implemented by Obedzinski et al. (2018).

### [B] Statistical Analysis

Wetted habitat data was compared with descriptive statistics as percent of wet, intermittent, and dry channel in the control and treatment reaches during 2019 and 2020 (Table 2, Group A). For most other variables, we used multiple before–after-control-impact timeseries (mBACIPs, Table 2, Groups B) or categorical and (mBACI, Table 2, Groups C) statistical analysis (Keough and Mapstone 1997, Wauchope et al. 2020). This follows our study design, which included multiple sites (riffle-pool units) in the *control* (not augmented) and *impact* (augmented) reaches, and sampling occurring *before* and *after* treatment (augmentation). In the mBACI experiments, the interaction of “Time” (before or after augmentation) and “Location” (control or impact reach) was considered significant when change occurred for the impact reach but not the control reach (e.g. Popescu et al., 2012; Smokorowski and Randal 2017). For time series data (Table 2, Group B) we constructed mBACI time series analysis following methods of Wauchope et al. (2020, EQ 1):

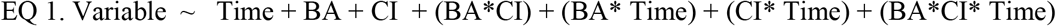

where time is a daily timestep centered on the day of augmentation; BA is a categorical variable of either “before” or “after”; and CI is a categorical variable of either “control” or “impact.”

**Table 2.** Statistical design used to evaluate each response variable by group type in the flow augmentation experiment.

| Group | Variable | Type | Analysis |
| --- | --- | --- | --- |
| A | Wetted Habitat | Categorical (manual measurement) | Summary Statistics |
| B | Water Temperature<br>Dissolved Oxygen<br>RCT Depth<br>Fish movement | Continuous logger or antenna detections | mBACIPs linear mixed effects model.<br>Variable ~ treatment* reach*time + (1\|site) + (1\|date) |
| C | Pool Velocity<br>Pool Depth<br>BMI Drift | Categorical (manual measurement) | mBACI linear mixed effects model.<br>Variable ~ treatment* reach + (1\|site) |
| D | Fish Growth | Categorical (capture mark recapture) | Control vs Impact ANOVA. |
| E | Fish Survival (by species) | Categorical (capture mark recapture) | MARK analysis.*<br>1. Null<br>2. Treatment<br>3. Year<br>4. Treatment and Year |

\*Models were run separately for each species (Coho Salmon or steelhead).

For categorical mBACI data (Table 2, Group C) we used the same model except without the continuous “Time” variable. Site (riffle-pool unit) was added as a random effect in all models. For both timeseries and categorical data (Table 2, Groups B and C) we used linear mixed-effect models with the LMER command in R version 3.5.1. To determine an augmentation treatment effect, we used the interaction BA*CI as the response variable (Table 2, Group C) and to determine an augmentation trend effect we used the interaction Time*BA*CI (Table 2, Group C), Wauchope et al. (2020).

We could not evaluate the effect of augmentation on juvenile salmonid growth using a BACI design due to insufficient sampling intervals. Reach-scale differences on growth were therefore described using a control impact (CI) analysis with ANOVA and discussed qualitatively (Table 2, Group D). For survival analysis (Table 2, Group E), we developed robust design capture-mark-recapture models using the PIT-tag wand data as described by Obedzinski et al. (2018) and used model evaluation to determine whether survival probability differed between control and impact reaches before and after augmentation. For each species, we constructed models in Program MARK (White and Burnham 1997) that allowed survival to vary by treatment (control or impact), time (before and after augmentation), and year (2019 or 2020). We also included a null model that held survival constant across treatment and year. In all models, detection probability was allowed to vary by reach and survey. Akaike’s Information Criterion corrected for small sample size (AICc) was used to evaluate model support, and models were considered to show similar support if they were within 0-2 AICc units and/or carried >10% of total model weight (Burnham and Anderson 2002). For the models with highest support, we then estimated the effect size using methods described in Cooch and White (2019).

## [A] Results

### [B] Wetted Habitat

The effects of augmentation on wetted habitat and surface flow varied between years but also expressed consistent spatial patterns. Stream drying occurred in low-gradient reaches, characterized by alluvial gravel deposits and wide bar-riffle morphology, especially downstream of rkm 0.5 and also between rkm 1.75 and 4.0 in the drier summer of 2020 (Figure 4). These reaches were previously identified as being particularly susceptible to drying (California Sea Grant, 2019, Supplemental Materials Figure S3). In 2019, after a May freshet, Porter Creek maintained surface connectivity throughout the summer except in the lowest reaches closest to the confluence with the Russian River. It is unclear whether augmentation effected channel connectivity during our study period in 2019; however, by October, 30% of the control reach was dry or intermittent, whereas only 12% of the impact reach was dry or intermittent (Supplemental Materials Table S1). Augmentation had a much larger effect on wetted habitat during the dry summer of 2020. Prior to augmentation (June 24, 2020), 92% of the control reach was still wetted, whereas only 56% of the impact reach was wet. One month after augmentation (July 30, 2020), the control reach experienced significant drying and wetted habitat decreased from 92% to 50%, whereas, wetted habitat in the impact reach increased from 56% to 90% (Figure 4, Supplemental Materials Table S1).

**Figure 4.**
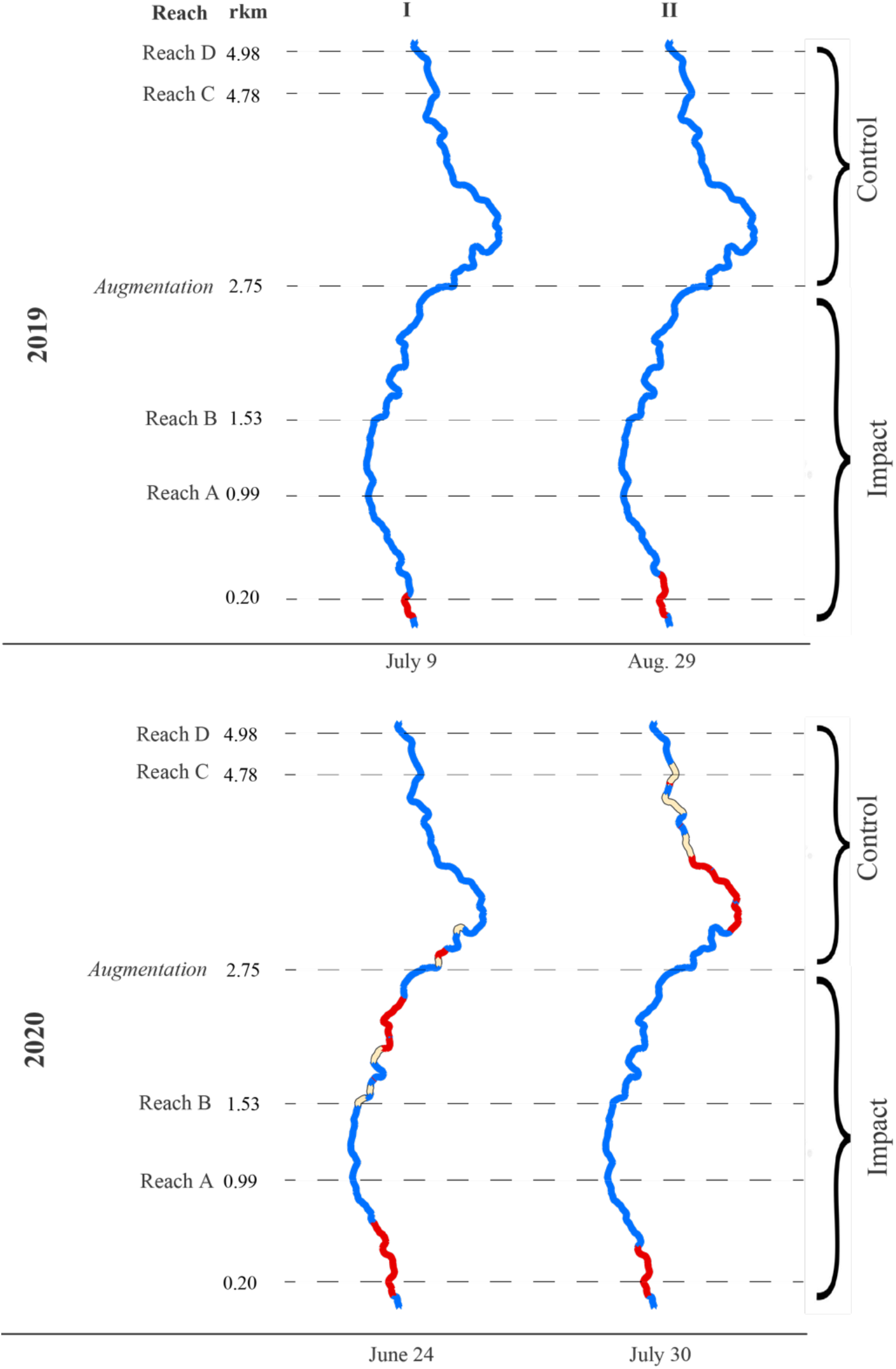
Wetted habitat maps showing flowing (blue), intermittent (yellow), and dry (red) stream channel immediately before (I), and following one month (II) of a 13.9 L/s augmentation in 2019 (top) and (2020) bottom, at sites upstream and downstream of the augmentation at rkm 2.75. The start (downstream end) of each study reach is shown with a dashed line.

### [B] Water Quality

Augmentation significantly increased dissolved oxygen in the impact sites during both years, although the effect size was much larger during the dry summer of 2020 (2.26 mg/L) than the wet summer of 2019 (0.23 mg/L) and the trend was only significant in 2020 (0.15 mg/L/day) (Table 3). During the wet summer (2019), augmentation had a small cooling effect on daily average water temperature (−0.41 C) and a negligible, (but statistically significant) effect on the trend (+0.043 C/day). During the dry summer, augmentation increased water temperature (+1.34 C), although water temperature stayed within suitable ranges for rearing salmonids. There was no effect on the trend. Figure 5 illustrates the effect of augmentation on water temperature, dissolved oxygen, and riffle crest depth at a single site in the impact reach.

**Table 3.**
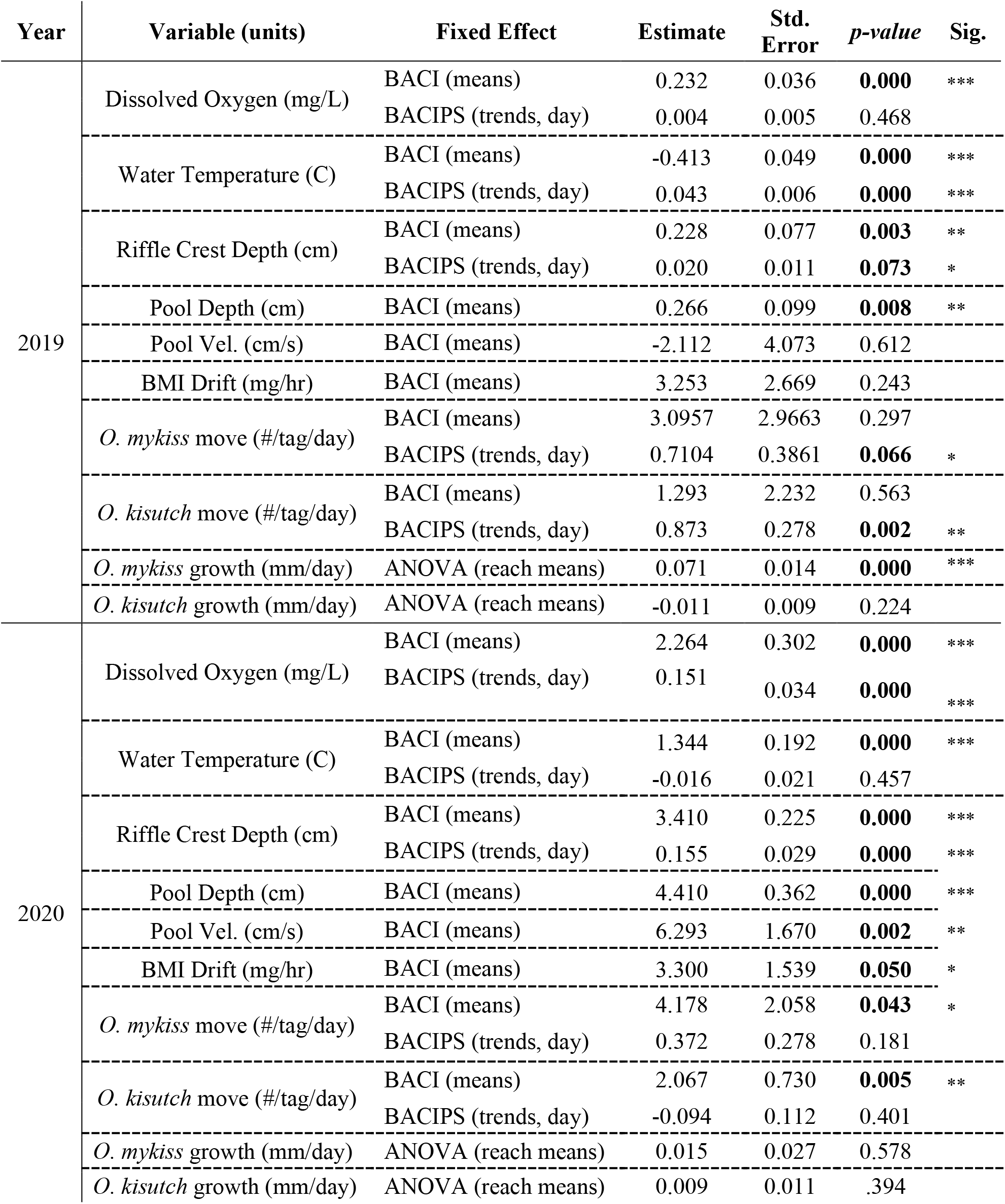
Results from mixed modeling for BACI (comparison of means) and BACIPs (comparison of trends) effects from augmentation in 2019 and 2020.

**Figure 5.**
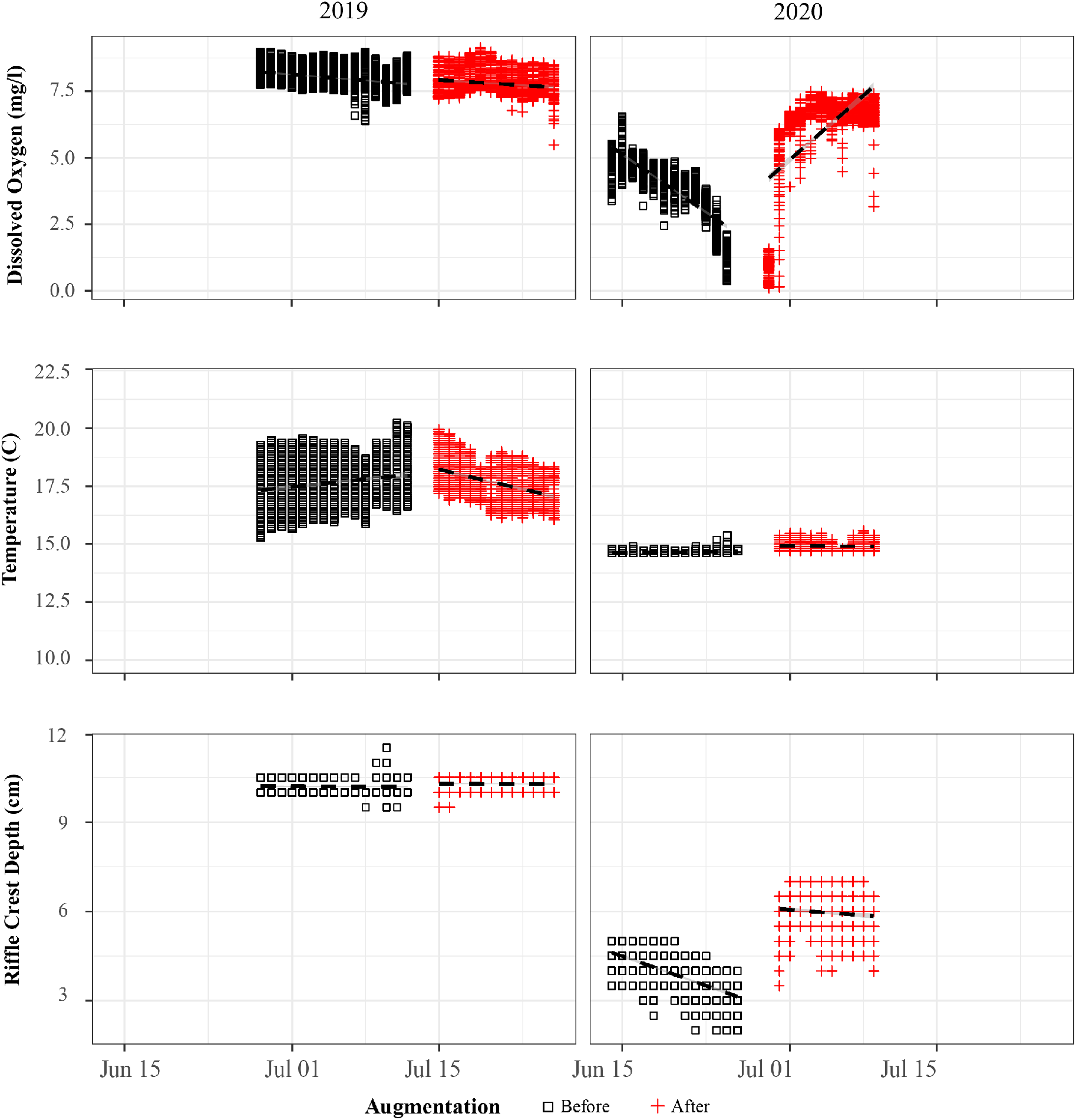
Dissolved oxygen (top), water temperature (middle),), and riffle crest depth (bottom), shown for representative impact site in reach B (Figure 2) to illustrate the before (blue) and after (red) periods in both 2019 (left) and 2020 (right).

### [B] Hydraulic Habitat

During the wet summer of 2019, augmentation had no effect on pool velocity, and caused a small, but statistically significant increase on riffle crest depth (+0.23 cm mean and +0.02 cm/day trend) and pool depths (+0.27 cm mean) (Table 3). During the dry summer of 2020 augmentation had a much larger effect on riffle crest depth (+3.4 cm increase in RCT depth and +0.155 cm/day trend) and pool depth (+4.4 cm mean). Pool velocity also increased significantly (+6.3 cm/s) due to augmentation in the dry summer (Table 3).

### [B] Stream Invertebrates

Although drifting invertebrates were near annual minimums by mid-summer in Porter Creek (Rossi et al. 2022), drift rate in the impact reaches declined less or increased relative to the control reaches following augmentation in both 2019 and 2020 (Figure 6); however, the effect of augmentation on invertebrate flux was only significant in 2020 (Table 3). Following augmentation in 2020, drifting invertebrate flux increased in the impact reach by 3.3 mg/hr relative to the control site (Table 3). The biomass flux of drifting invertebrates (mg/hr) was 66% to 85% higher in impact sites after augmentation than before, although the total number of drifting invertebrates declined slightly (Supplemental Materials Figure S4). In 2020 the impact reach saw increases in *Chironomidae* (midges) and *Baetids* (mayflies) after augmentation, which are vulnerable to predation by juvenile salmonids (Supplemental Materials Figure S4). The mean biomass of drifting invertebrates was nearly identical between control and impact reaches prior to augmentation, but mean biomass decreased more in the control reaches after augmentation (Supplemental Materials Table S2).

**Figure 6.**
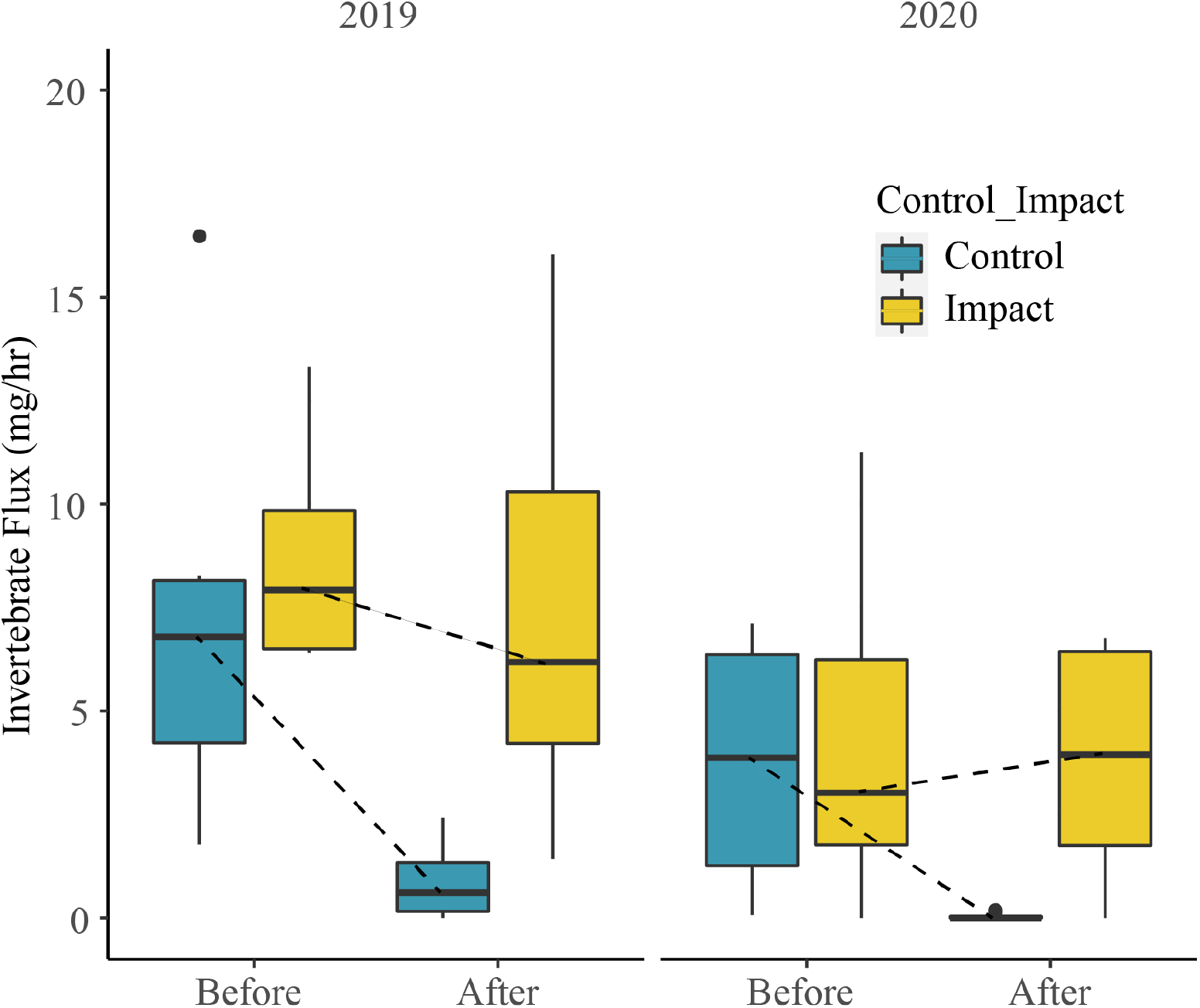
Mid-summer benthic invertebrate drift (mg/hr) before and after the onset of flow augmentation in Porter Creek during the control (blue) and treatment (yellow) reaches during in a wet year (2019) and a dry year (2020).

### [B] Salmonid Movement

In the wet summer of 2019, augmentation had no significant effect on total detections per tag of salmonids at riffle antennas, but the rate of detections per tag per day (trend) showed a significant increase after augmentation for both steelhead (0.71 detections/tag/day) and coho salmon (0.87 detections/tag/day) (Table 3). During the dry summer of 2020, augmentation significantly increased the total detections per tag of juvenile steelhead (by 4.2 detections per tag) and coho salmon (2.1 detections per tag) but had no significant effect on trend for either species (Table 3).

### [B] Salmonid Growth

The effect of augmentation on salmonid growth cannot be assessed using the BACI design since we did not have two growth intervals (before and after) in each reach (control and impact). During both years, the growth interval spanned the “before” and “after” augmentation periods with at least 30 days of growth after augmentation. However, an ANOVA analysis at the reach scale we found a small but significant effect on growth for young-of-year steelhead during the wet summer of 2019, when steelhead in the impact reach grew 0.071mm/day more than those in the control reach (Table 3). No effect on steelhead growth occurred in the dry year, and no effect on Coho growth was observed in either year.

### [B] Salmonid Survival

In the wet summer of 2019, the probability of survival for Coho Salmon increased in the impact reach during the interval in which the flow augmentation occurred (Figure 7). While the probability of survival for steelhead declined in both the control and impact reaches in 2019, the decline was significant only in the control reach. In 2020, the dry year, we observed a decline in survival probability from the pre-augmentation interval to the post-augmentation interval in all reaches; however, for both species the magnitude of the decline was greater in the control reaches. For juvenile Coho Salmon, the effect of rearing in the augmentation reach was an increase in survival probability of 0.11 in 2019 (wet year) and 0.24 in 2020 (dry year) (Figure 8). For steelhead, there was no significant effect of rearing in the augmentation reach in 2019, and in 2020, it increased survival probability by 0.20.

**Figure 7.**
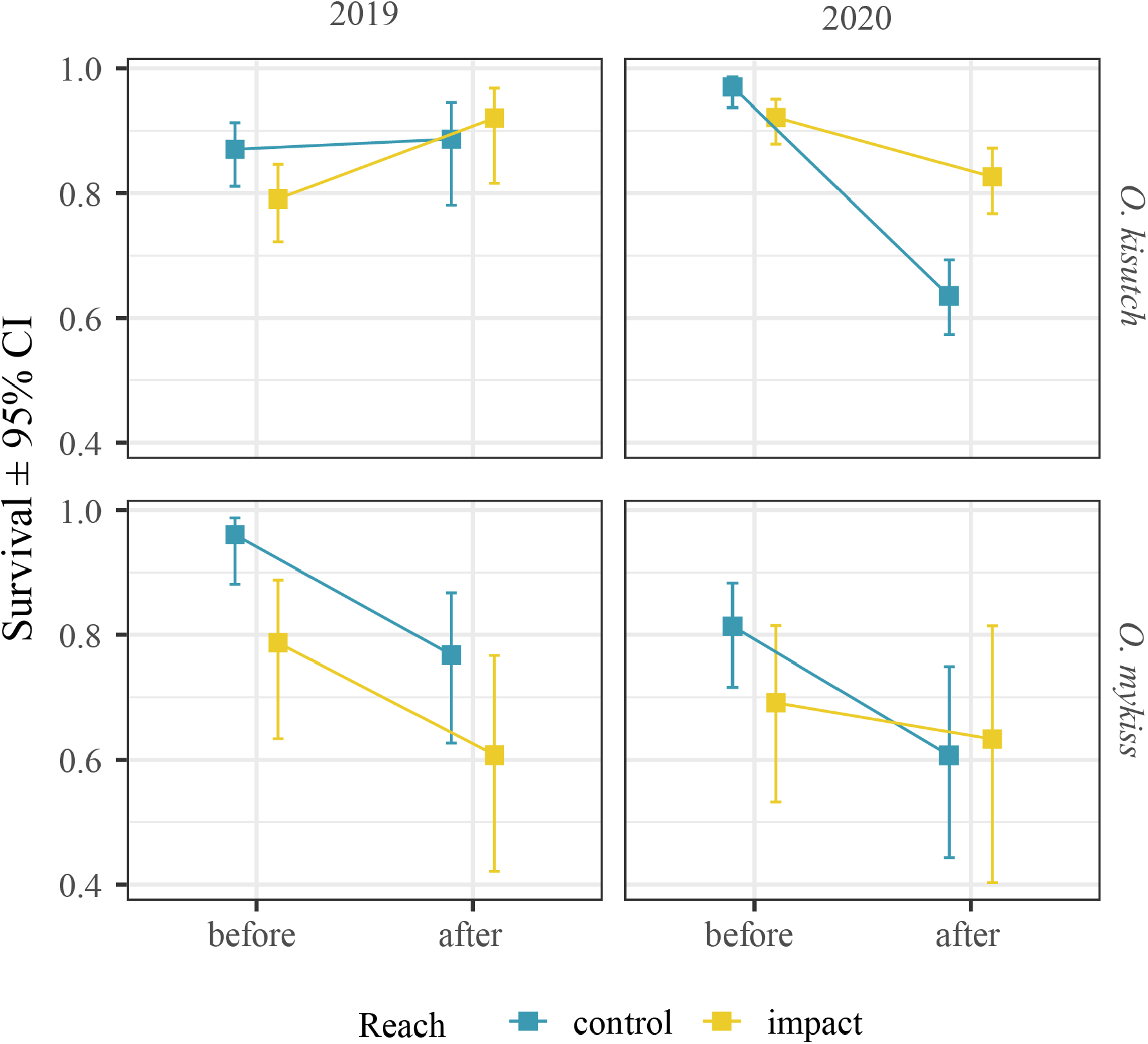
Probability of juvenile salmonid summer survival before and after the onset of flow augmentation in Porter Creek in a wet year (2019) and a dry year (2020). Change in probability from control reaches is shown as the blue line, and for impact reaches as the yellow line.

**Figure 8.**
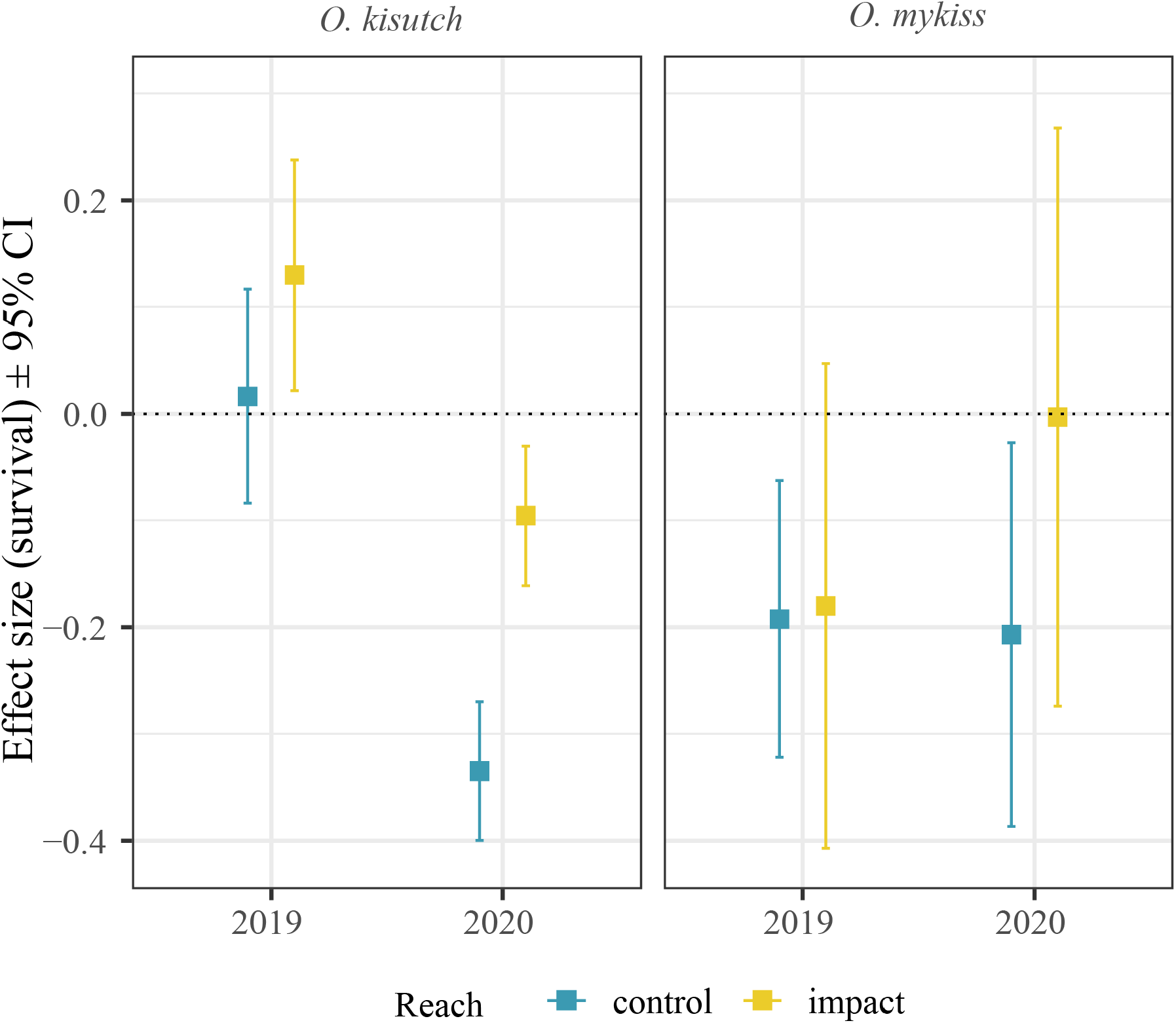
Effect size of juvenile salmonid survival before and after flow augmentation in control and impact reaches in a wet year (2019) and dry year (2020).

## [A] Discussion

There are numerous studies that illustrate the negative ecological effects of streamflow impairment, which have primarily focused on large, regulated, or dammed rivers (Carlisle et al. 2019). While it is self-evident that flow-impaired streams should benefit from augmentation, few studies have quantified the ecological responses to flow restoration in small, undammed streams (Davies et al. 2014, Gillespie et al. 2015). To mitigate flow impairment in undammed streams there is growing interest in novel projects including groundwater management (Woelfle-Erskine et al. 2017) and the direct release of water into streams from off-stream storage (Deitch and Dolman 2017, Ruiz et al. 2019), as illustrated in this study. But question remain over the magnitude, timing, and duration of flow restoration required to produce a meaningful ecological benefit. Our study provides evidence that a well-timed, short-duration, low-volume, flow augmentation measurably improved the habitat quality and survival of summer-rearing salmonids. It suggests that, in systems like Porter Creek, even slight increases in streamflow (e.g. 10-20 L/s) during critical periods can move salmonids away from the knife’s edge of survival and potentially aid in the recovery of imperiled populations. Our results and discussion focus on the flow augmentation case study; however, we suggest they can also serve the broader project of environmental flow management as empirical evidence for the benefits of small-scale flow restoration.

### [B] Spatial and Temporal Effects of Augmentation on Habitat Connectivity

Flow augmentation had a direct positive effect on hydraulic connectivity, with the influence diminishing downstream, consistent with our hypotheses (H1 and H2). However, we also found that the effects of summer augmentation on hydraulic connectivity (and all variables) were more sensitive to ambient streamflow than to distance downstream from the point of augmentation. In 2019, ambient flow remained relatively high and we observed negligible effects on downstream streamflow or connectivity from augmentation treatments during the study period (Figure 4). In 2020, significant drying occurred prior to augmentation, especially in reaches with deep, porous gravel deposits (around rkm 2 and downstream of rkm 0.6), which is consistent with findings from research in coastal California. For example, Lovill et al. (2018) found that sites with conductive, in-channel sediment fill were most likely to experience surface flow disconnection during low flows in nearby Elder Creek. Moidu et al. (2021) found that drying in Russian River tributaries tends to occur in low gradient alluvial reaches, which often characterize the downstream extent of tributary streams in the Russian River. In our study, one month of augmentation at 13.9 L/s was able to re-wet most of the downstream channel in Porter Creek in 2020; however, it is unclear whether continuing this level of augmentation would have maintained the wetted channel through the rest of the fall season. Nonetheless, reduced duration of stream disconnectivity is strongly related increased survival of salmonids, as has been documented by others (Obedzinksi et al. 2018; Vander Vorst 2020, Hwan and Carlson 2016; Sloat and Osterback 2012; Woelfle-Erskine et al. 2017).

### [B] Augmentation Effects on Water Quality and Hydraulic Habitat

We found that flow augmentation increased dissolved oxygen, in support of our hypothesis (H3). Dissolved oxygen increased significantly in both years (Table 3); however, the effect size was negligible in the wet summer of 2019 and had much greater ecological significance in 2020. In the two weeks prior to the 2020 augmentation, DO in the BACI study impact sites (median of 4.2 mg/L, SD 2.6 mg/L) had lowered to levels known to impair swimming performance and food conversion efficiency for juvenile Coho Salmon and steelhead (Bjornn and Reiser 1991, USEPA 1986). Augmentation increased dissolved oxygen to a median of 6.6 mg/L (SE 1.52 mg/L), at which juvenile salmonids experience minimal impairment (USEPA 1986).

We found marginal support in Porter Creek for our hypothesis (H4) that augmentation would influence water temperature. In this study, augmentation had a statistically significant but relatively small effect on water temperature – slightly decreasing temperature in the cooler summer of 2019 (−0.43 C) and increasing temperature in the warmer summer of 2020 (+1.3 C). Since augmentation resulted in a small change in the total volume of water in the channel, we suspect that atmospheric and within-channel transport processes largely controlled stream water temperature downstream from the point of augmentation and buffered the effects of warmer augmentation at downstream study sites. Temperatures at all our sites remained below stressful physiological tolerance limits of salmonids. However, we acknowledge the potential for augmentation to significantly affect stream temperatures under different site conditions, including initial temperatures (from the pond), flow release rates, flow of receiving water bodies, and other local environmental factors (Ficklin et al. 2012).

While the effect of augmentation on riffle and pool depths was significant in both summers, the effect size was much larger in the dry summer of 2020. The increase in riffle depths in 2020 were equivalent or greater than common body depths of steelhead and Coho smolts (Negus 2003) and co-occurred with increased detections of steelhead at riffles (Table 3). Riffle depths over 4 cm corresponded to an increase in detections of juvenile steelhead in riffles during 2020. More work is needed to understand the mechanisms behind this relationship.

### [B] Augmentation Effects on Stream Invertebrate Drift

The biomass of invertebrate drift increased relative to control reaches following the onset of augmentation, consistent with our hypothesis (H5) and previous studies, which have shown that increased wetted area and riffle velocity can increase the production, hydraulic transport, and behavioral drift of invertebrate species (Annear et al. 2004, Svendsen et al. 2004; Naman et al. 2016). However, the natural phenology of invertebrate drift in Porter Creek was near its annual minimum by during mid-summer (Rossi et al. 2022) and the relative increase in drift of less than 4 mg/hr represents a small change in the growth potential of foraging salmonids. For example, Porter Creek drift rates in April, 2018 were between 100 and 200 mg/hr, excluding large or rare invertebrates (Rossi 2022). Using published energy densities for freshwater invertebrates (e.g. 3,072 joules/gram Thompson and Beauchamp 2016), an additional 4 mg/hr would only add a negligible ~12.3 joules per hour of potential energy for salmonids. Late-summer growth rates for salmonids are naturally near zero in many California coastal streams (Kelson et al. 2019, Rossi et al. 2022). Thus, we suspect that the benefits to growth and survival from this level of augmentation (13.9 L/s) in mid-summer are more likely due to decreased metabolic stress from increased dissolved oxygen, or perhaps increased mobility of fish, than this modestly increased drift rate.

### [B] Augmentation Effects on and Salmonid Movement, Growth and Survival

As predicted (H6), augmentation was associated with increased detections per tag in both years, but the results were only significant in the dry summer of 2020. This was not surprising since hydraulic connectivity and suitable dissolved oxygen concentrations were largely maintained in 2019, whereas the increase in riffle depth and dissolved oxygen following augmentation in 2020 significantly improved the hydraulic and metabolic environment for fish movement. We chose the ‘number of detections per tag’ as our response variable to indicate inter-pool movement, since we observed that some animals were detected much more frequently than others. A further analysis of covariates with detections (e.g. dissolved oxygen, riffle crest depth, and velocity) would greatly aid our understanding of the mechanisms by which streamflow augmentation affected salmonid movement. Since PIT antennas were placed over riffle habitats and away from pools, and since riffles had become very shallow by late summer, we reasoned that most detections were associated with movement between pools; however, we cannot rule out that some detections were related to foraging movements into riffles and not movement between pools.

In our evaluation of growth in control versus impact reaches, we observed a statistically significant effect for young-of-year *O. mykiss* in 2019 (0.071 mm/day higher in the treatment reach, Table 2), but no effect in 20202 of for *O. kisutch* in either year. While factors that we did not account for in this study may have been influencing growth, such as density-dependence and inter-specific interactions, it is reasonable to infer that increases in dissolved oxygen and riffle depth from augmentation aided fish growth. However, our sample design, with growth intervals before and after augmentation, was not suited for to statistically test the effects of augmentation on growth. Exploring the effect of augmentation on foraging behavior (Rossi 2021b) and evaluating changes to drift and water quality in a bioenergetic model could help to elucidate the mechanisms by which augmentation may affect the growth of salmonids.

Perhaps the most consequential finding of this study was the significant increase in over-summering survival (H8) related to the flow augmentation for both juvenile Coho Salmon (+24%) and steelhead (+20%) in 2020 and for juvenile Coho Salmon (+11%) in 2019. Previous work on salmonid over-summering mortality in Russian River tributaries indicated that days of stream disconnectivity (intermittent flow) and low dissolved oxygen had strong negative correlations with Coho Salmon over-summer survival (Obedzinski et al. 2018), whereas pool volume and water temperature were weakly correlated with survival. Our study suggests that augmentation, particularly in the dry summer of 2020, strongly effected those variables most closely associated with salmonid survival (i.e., stream connectivity and dissolved oxygen), although this relationship may not be transferrable to other systems.

### [B] Future Work

While the findings of increased over-summer survival of coho and steelhead flow augmentation are promising, more work is needed to understand the life-cycle consequences of these effects (e.g. at the population level) and their dependencies on ambient environmental conditions. Although 2020 was a dry year, it is uncertain that this level of flow augmentation would confer the same benefits in critically dry years or how the physical and biological effects we measured scale with augmentation rates. The ability to implement flow augmentation in other systems will also depend on several factors, including the availability of relatively large storage systems with reliable water supplies, infrastructure to manage and monitor flow releases, willing landowners, and the support of natural resource agencies. Finally, more work is needed to monitor effects of augmentation on non-target species (e.g. stream amphibians and non-salmonid fishes), understand the capacity of augmentation to affect adult salmon survival, and investigate the ecological effects of altering the duration of connectivity in an intermittent stream over many years. Nonetheless, given the critical state of salmonid populations in California (Moyle et al. 2017), the increased development of small-scale water storage in the study region (Dietch et al. 2013), and new models of collaboration between agriculture and wildlife conservation (Holmes et al. 2021), the findings of this study suggest that flow augmentation may be an important tool within the broader salmonid conservation strategy.

## [A] Conclusion

To our knowledge this study represents the first controlled flow manipulation experiment to quantify the effects of flow augmentation from off-channel storage on juvenile salmonids. Numerous studies illustrate the negative ecological effects of flow impairment but few if any of experimental studies have quantified the consequences of flow restoration. To date, flow augmentation projects have been implemented opportunistically as a novel restoration strategy for salmonids in Northern California (Deitch and Dolman 2017, Ruiz et al. 2019). Our findings suggest that appropriately timed and located flow enhancement may significantly improve habitat connectivity, the metabolic environment, and survival of summer rearing salmonids. More broadly, this project provides a template for how to monitor the effects of flow enhancement in small streams, highlights critical areas for further study, and provides empirical evidence that environmental flow programs which effectively restore streamflow in small, salmon-bearing streams can yield significant ecological benefits.

## [A] Acknowledgments

Critical field and analytical support for this project came from California Sea Grant staff, especially Andrew Bartshire and Elizabeth Ruiz. We also acknowledge field and laboratory support from UC Berkeley students Gunnar Reith, Sam Larkin, Shannon Mckillop-Herr, Alia Aguilar, Michael T Scheweiker, and Nora Povejsil. Stream gauging was conducted by Trout Unlimited staff led by Mia van Docto and development and coordination of the augmentation project was led by Katie Robbins. We thank Gallo Vineyards, especially John Nagle and Tyler Hammond, for access and continued support for our scientific research, as well as their willingness to release water into the stream for the benefit of fish. Support from California Department of Fish and Wildlife staff David Hines and Corinne Gray was also much appreciated. Project design and construction was funded through private, state, and federal funds, including a 2015 Wildlife Conservation Board (WCB) grant (Project WC-1553M) and a Junior Investigator Award from the California Institute of Water Resources (Project 2016CA370B). Gabriel Rossi was additionally supported by a National Science Foundation CZP EAR-1331940 for the Eel River Critical Zone Observatory.

## [A] Data and Code Availability

https://datadryad.org/stash/share/SNsDJsHRQCAtPv0QGJnCbhLK6fY5W9ZnbmfOFPTwPlQ

## Supplemental Materials

**Table S1.** Summary Statistics of Wetted Channel Mapping in 2019 and 2020.

| Location | Date | Augmentation | Condition | Proportion of Reach |
| --- | --- | --- | --- | --- |
| control | 7/9/19 | Before | Wet | 100% |
| impact | 7/9/19 | Before | Dry | 1% |
| impact | 7/9/19 | Before | Wet | 99% |
| control | 8/29/19 | During | Wet | 100% |
| impact | 8/29/19 | During | Dry | 7% |
| impact | 8/29/19 | During | Wet | 93% |
| control | 10/11/19 | During* | Dry | 13% |
| control | 10/11/19 | During* | Intermittent | 16% |
| control | 10/11/19 | During* | Wet | 70% |
| impact | 10/11/19 | During* | Dry | 12% |
| impact | 10/11/19 | During* | Intermittent | 1% |
| impact | 10/11/19 | During* | Wet | 87% |
| control | 6/24/20 | Before | Dry | 3% |
| control | 6/24/20 | Before | Intermittent | 5% |
| control | 6/24/20 | Before | Wet | 92% |
| impact | 6/24/20 | Before | Dry | 35% |
| impact | 6/24/20 | Before | Intermittent | 9% |
| impact | 6/24/20 | Before | Wet | 56% |
| control | 7/30/20 | During | Dry | 27% |
| control | 7/30/20 | During | Intermittent | 23% |
| control | 7/30/20 | During | Wet | 50% |
| impact | 7/30/20 | During | Dry | 10% |
| impact | 7/30/20 | During | Wet | 90% |
| control | 10/01/20 | After | Dry | 99% |
| control | 10/01/20 | After | Wet | 1% |
| impact | 10/01/20 | After | Dry | 69% |
| impact | 10/01/20 | After | Intermittent | 9% |
| impact | 10/01/20 | After | Wet | 23% |

**\*Note in October 2019 augmentation was continuing, but rates had dropped below 13.9L/s due to head loss in the reservoir.**

**Table S2.**
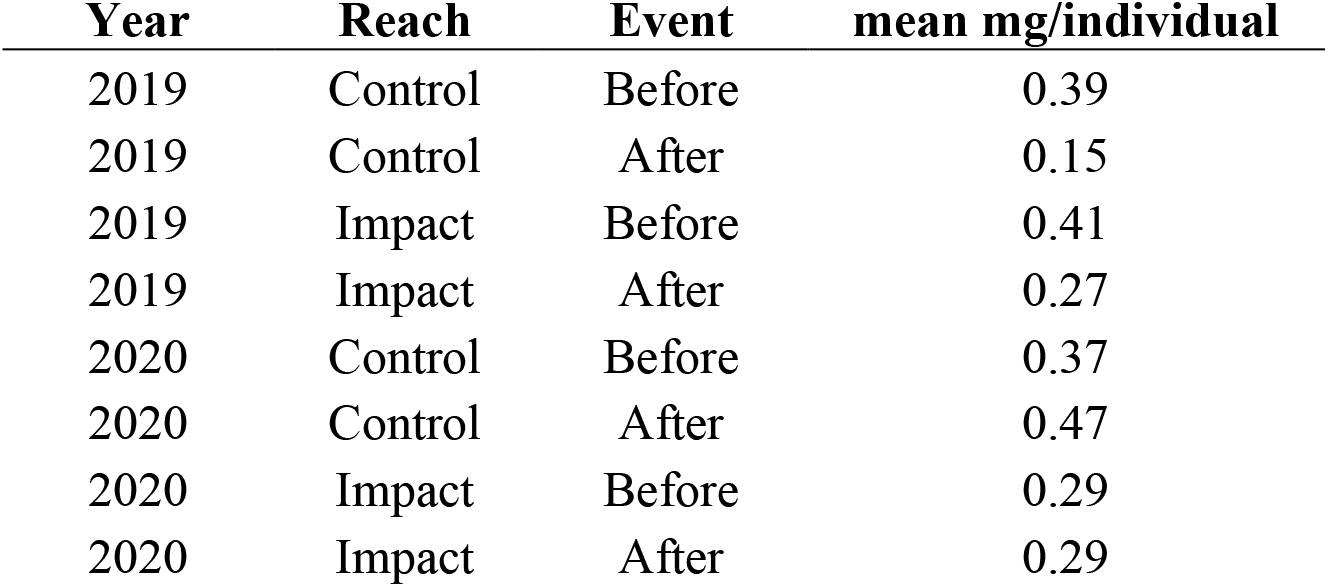
Mean biomass per individual invertebrate before and after augmentation, in control and impact reaches during 2019 and 2020.

| **Year** | **Reach** | **Event** | **mean mg/individual** |
| --- | --- | --- | --- |
| 2019 | Control | Before | 0.39 |
| 2019 | Control | After | 0.15 |
| 2019 | Impact | Before | 0.41 |
| 2019 | Impact | After | 0.27 |
| 2020 | Control | Before | 0.37 |
| 2020 | Control | After | 0.47 |
| 2020 | Impact | Before | 0.29 |
| 2020 | Impact | After | 0.29 |

**Figure S1.**
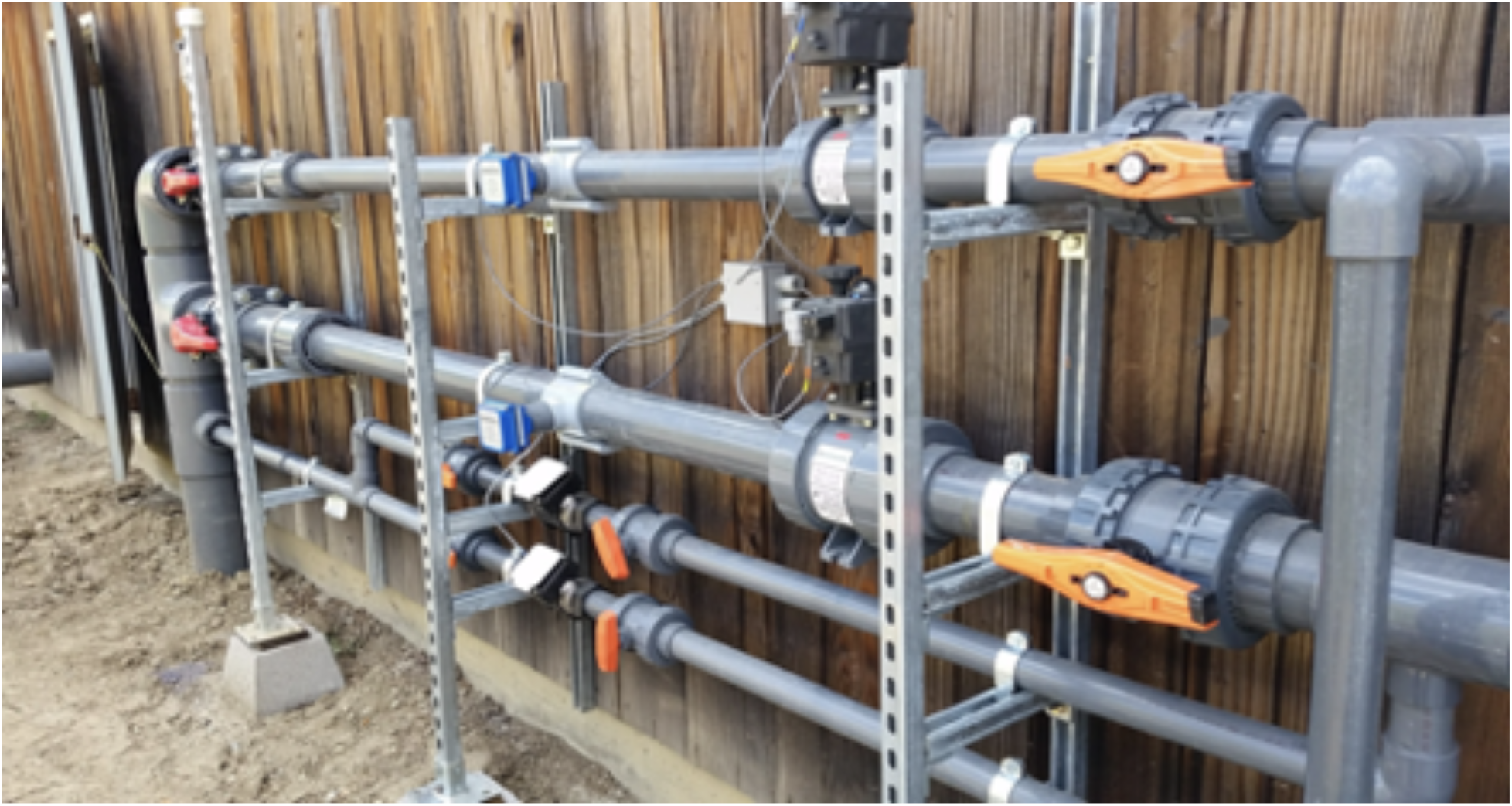
The plumbing manifold for the Porter Creek augmentation structure.

**Figure S2.**
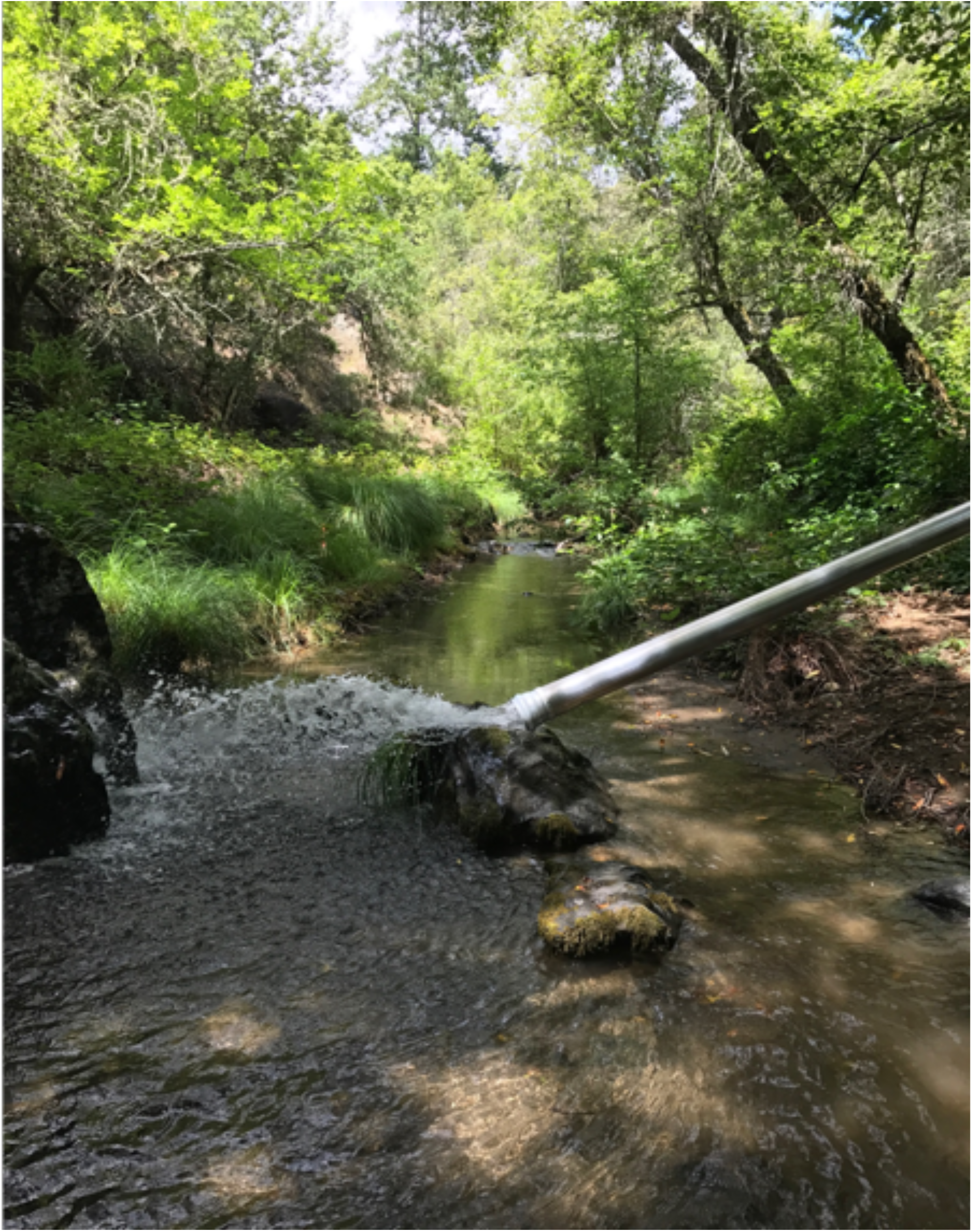
The lower augmentation point (see Figure 1) in Porter Creek, augmentation rate is 13.9 L/s.

**Figure S3.**
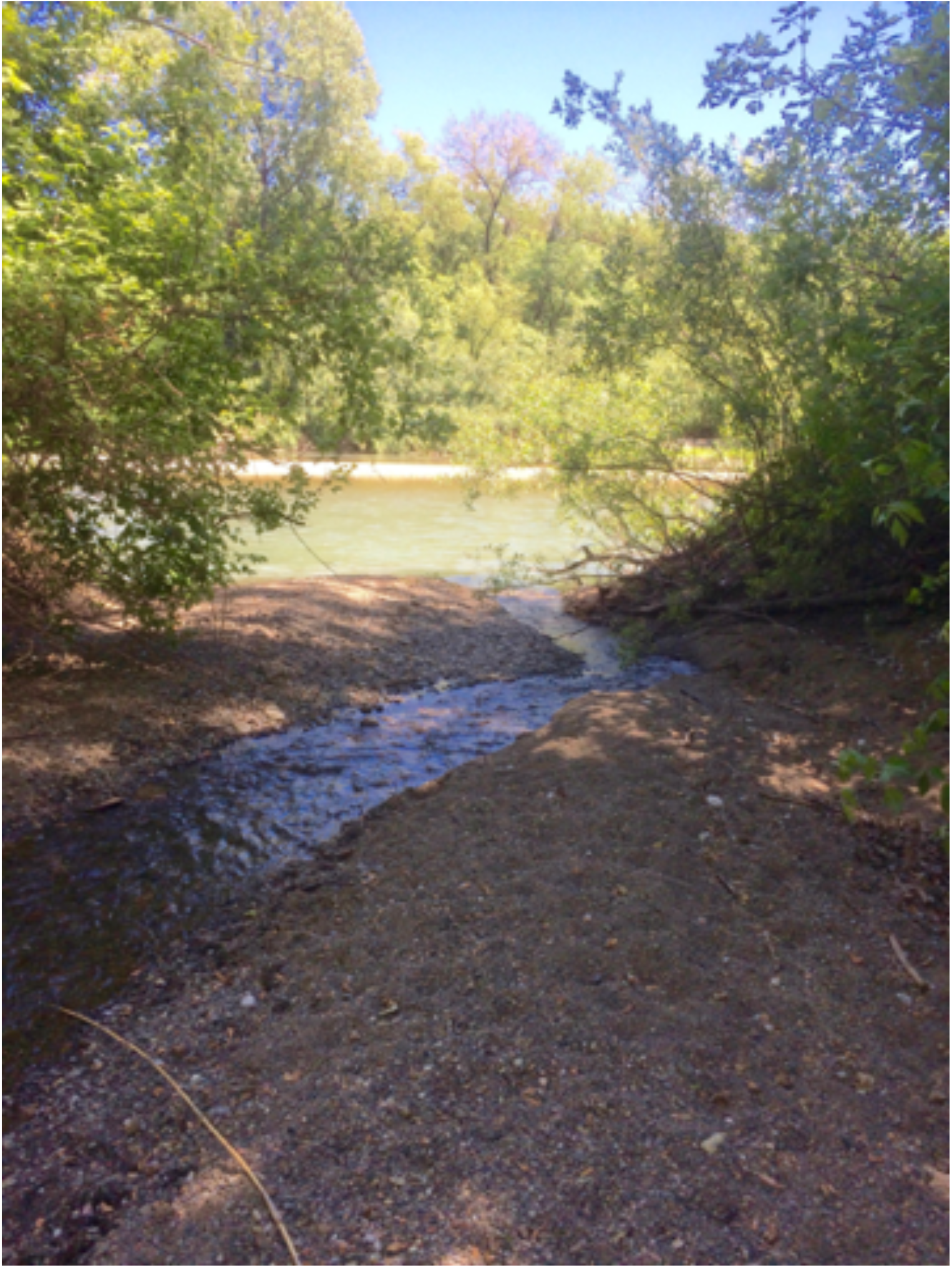
The Porter Creek – Russian River confluence showing the highly porous, and often backwatered channel of lower Porter Creek. This reach goes dry every year and seems particularly sensitive to river stage.

**Figure S4.**
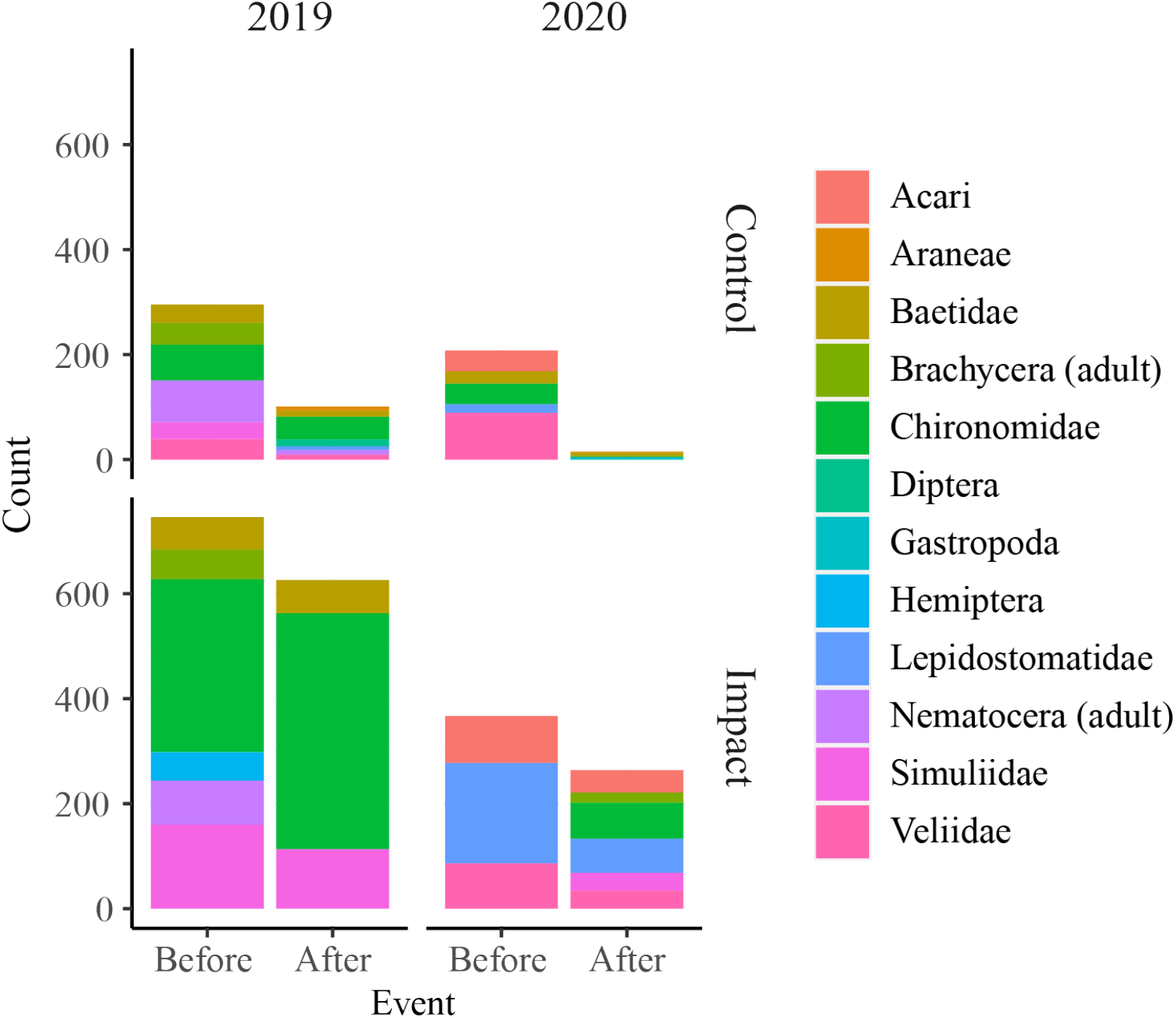
Counts of the twelve most common taxa found in drift samples, in 2019 (left) and 2020 (right), in the control (top row) and impact (bottom row) reaches, before and after augmentation.

